# *In vivo* capture of bacterial cells by remote guiding

**DOI:** 10.1101/2021.08.06.455395

**Authors:** Iaroslav Rybkin, Sergey Pinyaev, Olga Sindeeva, Sergey German, Maja Koblar, Nikolay Pyataev, Miran Čeh, Dmitry Gorin, Gleb Sukhorukov, Aleš Lapanje

## Abstract

Recently, it has been shown that several bacterial strains can be very efficient in cancer treatment since they possess many important properties such as self-targeting, ease of detection, sensing and toxicity against tumors. However, there are only a few relevant “candidates” for such an approach, as targeting and detection one of the biggest challenges as well as there are many limitations in the use of genetic approaches. Here, it is proposed the solution that enables surface modification of alive bacterial cells without interfering with their genetic material and potentially reduces their toxic side effect. By the electrostatic interaction fluorescently labeled polyelectrolytes (PEs) and magnetite nanoparticles (NPs) were deposited on the bacterial cell surface to control the cell growth, distribution and detection of bacteria. According to the results obtained *in vivo*, by the magnet entrapment of the modified bacteria the local concentration of the cells was increased more than 5 times, keeping the high concentrations even when the magnet is removed. Since the PEs create a strong barrier, *in vitro* it was shown that the division time of the cells can be regulated for better immune presentation.

## 1. Introduction

Using a top-down engineering approach, the ideal cancer therapy can be envisioned as micro or nano programmable “robot devices” that (i) specifically target tumors preferentially by self-propelling, (ii) are selectively cytotoxic only to cancer cells and (iii) responsive to external signals, which can transfer information about the local environment and will enable remote guiding.^[1]^ Specific targeting is one of the most important properties since it would allow the use of toxic substances without systemic effects. In addition, self-propulsion independently to the blood flow enables penetration into tumor regions that are otherwise inaccessible to the passive systemic deliverable therapies. According to above mentioned criteria for ideal anticancer treatment, bacterial cells seem to share many of these characteristics and in fact, there are genera, which have been shown in laboratory experiments as well as in clinical tests (Phase 1)^[2]^ to preferentially accumulate in tumors and lyse cancerous cells, including *Salmonella*,^[3]^ *Escherichia*,^[4]^ *Clostridium*,^[5,6]^ and *Bifidobacterium*,^[7]^ *Caulobacter*,^[8]^ *Listeria*,^[9,10]^ *Proteus*,^[11]^ and *Streptococcus*.^[12]^ Although there are very encouraging reports of anticancer bacterial therapy treatment, the current approach is far from the ideal cancer therapy since (i) in many cases native bacteria must be genetically modified at least to ensure safe use, (ii) specificity of delivery must be increased and (iii) it is needed to remotely monitor distribution of the bacterial cells. Currently it is known that the cells used in bacterial therapy are not directed specifically to the tumorous mass, but the accumulation is a passive process where bacterial propagation is only enabled within the locally deprived immunity,^[13]^ in contrast to the other tissues where bacteria are suppressed by strong immune responses of the host.

In addition, bacteria and their cellular components are also extremely potent and strong immune stimulators with endo-and exotoxins where their systemic delivery increases the risk of sepsis as well as development of toxic syndrome, since the only available control of their propagation in such approaches is by using antibiotics.^[14]^ Due to the above mentioned drawbacks, the bacterial therapy cannot be extended to the wide range of bacteria although some might have very potent toxins, e.g. *Pseudomonas* strains,^[15]^ since they cannot be directed to the tumor cells or because of very low scores on the above mentioned characteristics of the ideal bacterial agent that could be used in the approach.

Having in mind these drawbacks, especially, to increase repertoire of bacteria that can be used in treatments by the specific delivery to the site, monitoring their distribution and controlling their growth, here we propose the robust solution that solves these drawbacks without introducing genetic manipulations methods.

Since the bacterial cells are negatively charged, they can act as a core for deposition of PE layers on their surfaces using a Layer-by-Layer (LbL) approach, to form a tailor-made core-shell structure. Deposition of the PE on the cell surface can add new modalities to the bacteria such as survivability in the harsh conditions or improved adhesive properties.^[16]^ In addition, it can bring new physiological activities as increased specific binding to cells, or it can expand the spectrum of action of the cells using modified PEs with fluorescent dyes, enzymes or ligands (e.g. antibodies),^[17–19]^ as well as potentially allow them to get detected in the body. For the remote targeting of the bacteria, inside the layers it can also be added different sorts of NPs, which could be used for the controlled delivery as shown in delivery systems using nano-and microparticles.^[20]^ It is expected that such shells on the one hand separate the surface of the cell from the immune system and on the other, due to the shell strength, they can control the division rate^[21]^ and therefore, invasiveness of the delivered bacterial growth. Accordingly, our main aim in the study was to test an idea to guide, detect and monitor distribution and growth of bacteria within the mammal body in order to make safer anticancer or perhaps, also other therapies using bacteria, as well as to broaden the spectra of possible use of different bacterial strains. In relation to that, by using the *in vivo* experiments on the mouse model more specific aims were: (i) to remotely guide bacterial cells by using magnetic field, (ii) monitoring the distribution of bacterial cells inside the mammal body and (iii) controlling division rate of the selected strain.

## 2. Results

The surface of the bacterial cells was efficiently functionalized by the method of electrostatic attachment of the PEs labeled with fluorescent dye and the following consequent deposition of the magnetic NPs (**Figure 1**A and **1B**). In order to prepare Scanning Electron Microscopy (SEM) images of bacteria coated with PE and magnetic NPs it was found that just air drying is one of the most convenient and relevant approaches, since the chemical fixatives interfered with the structure of the shells, causing many artefacts such as aggregation or release of the cellular compounds (data not shown). To confirm the presence of iron NPs on the surface of the coated cells, we applied a combination of SEM and energy-dispersive X-ray spectroscopy (EDS) techniques (**Supplementary Table 1**). As a result of the coating, the magnetite NPs adsorbed on the polycation-treated bacteria provided a dense layer on the entire cell surface (Figure 1C and 1D).

**Figure 1.**
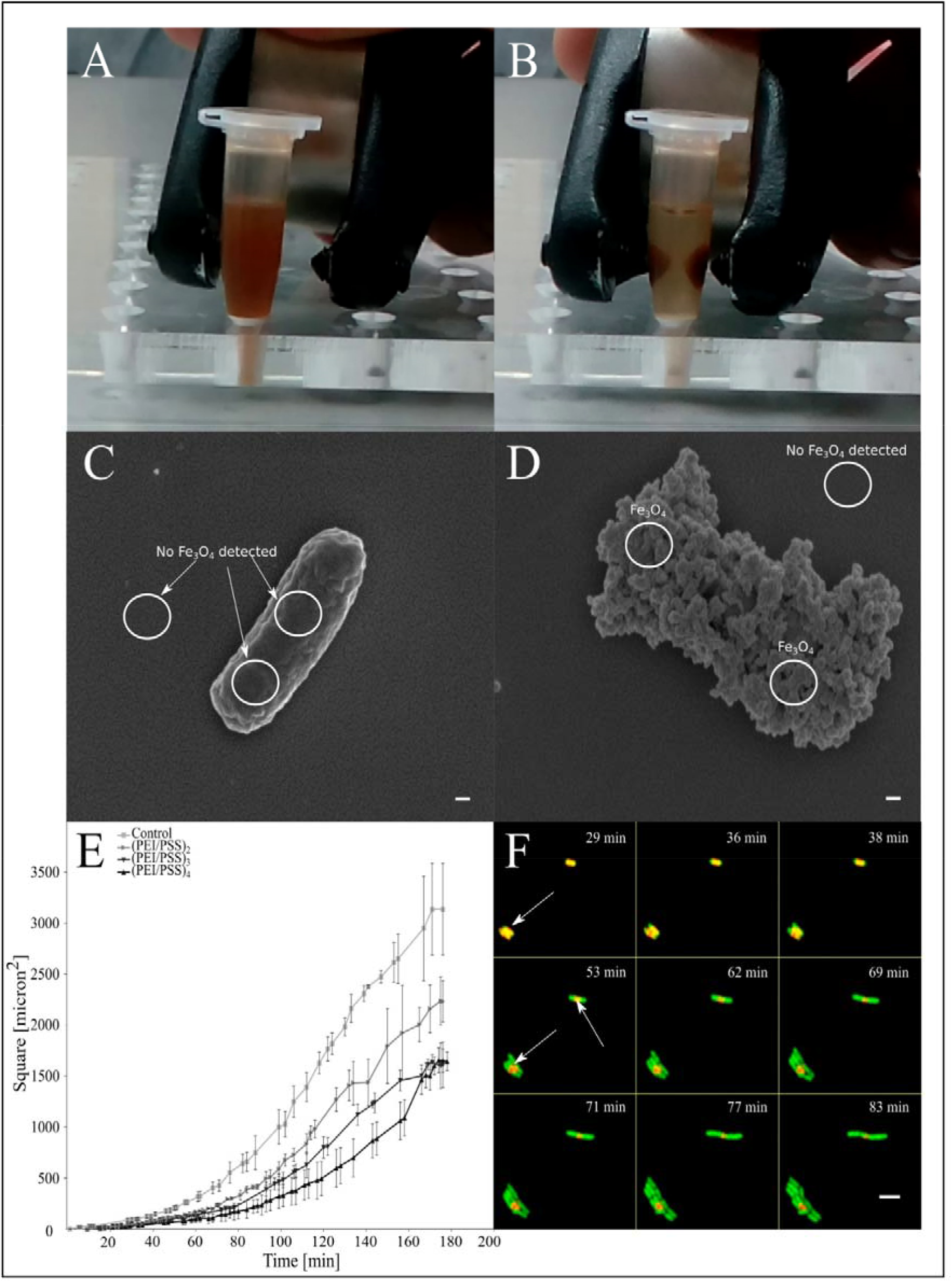
*In vitro* magnet entrapment of the *E. coli* cells coated by polyethylenimine(PEI)/poly(styrene)sulfonate (PSS)/PEI/Fe_3_O_4_ (A, B). When a magnet is applied, movement occurs instantly. Additionally, presence/absence of nanoparticles was proved by SEM (EDS), where intact cells did not have any particles, scale bar = 100 nm (C), while the modified cells had magnetite only on the surface (D). Effects of entrapment of cells in layers of polyelectrolytes on growth (E) of the population of the single bacterial cells. Escape of the cells from the LbL layers when entrapped in PE layers (F) red is LbL shell indicated with arrows, green is GFP producing cells, overlap regions are yellowish, scale bar = 5µm.

By time-lapse confocal microscopy (TLCM), we showed that the entrapped cells stay alive inside the formed PE layers. We determined that the most appropriate for the injection amount of layer was 4 bilayers deposited on the cells, which enabled us the highest possible fluorescence level, slower division rates of the cells and at the same time, keeping the aggregation of the cells during the LbL procedure as low as possible. After incubation *in vitro*, bacteria started escaping from the shells by breaking the capsule at the distal positions, leaving parts of PE capsule, which surrounded the escaping cells (Figure 1F). On the single cell level, we determined that the growth rate was proportional to the number of the PE layers (Figure 1E) deposited on the bacterial surface, where the slowest growth was observed for the cells entrapped in 8 layers. Although there was no toxicity, all the coated cells demonstrated on average 1.4 times longer lag phase than untreated cells. The cells coated in our procedure showed that they can be released from the capsules in 50 minutes after incubation in the most optimal conditions.

After administration of the electrostatically functionalized bacterial cells (EFBC) into the bloodstream of mice, we observed differences in distribution of cells that were dependent on the presence of the permanent magnet. Under exposure to the magnetic field, spreading of the cells in the organism from the place of injection was directed toward the paw with a magnet with the following accumulation of the bacterial cells, forming bright areas in the paw where permanent magnet was placed (**Figure 2** A) after a certain time. In contrast, within other parts of the body the signal became 2,4 times on average weaker than in the paw. In control groups, the EFBC were mainly distributed in the abdomen forming dense shining areas, whereas in the intact paw the signals were weaker 1.8 times than in other parts of the body (Figure 2 B). Throughout all experiments animals remained alive. By the magnet entrapment we were able to significantly increase concentrations of the cells more than 5 times in the paw region than in the control group (*P* < 0.001) (Figure 2C). However, although the magnet efficiently attracted bacterial cells, we determined that just a fraction of cells (up to 4.64%±0.14%) was accumulated in the paw, while the overall distribution of free circulating cells remained the same in both groups. After the magnet was removed, we observed that the signal from the exposed paw was 5.8 times higher than in unexposed paw over the remaining half an hour (*P* < 0.001).

**Figure 2.**
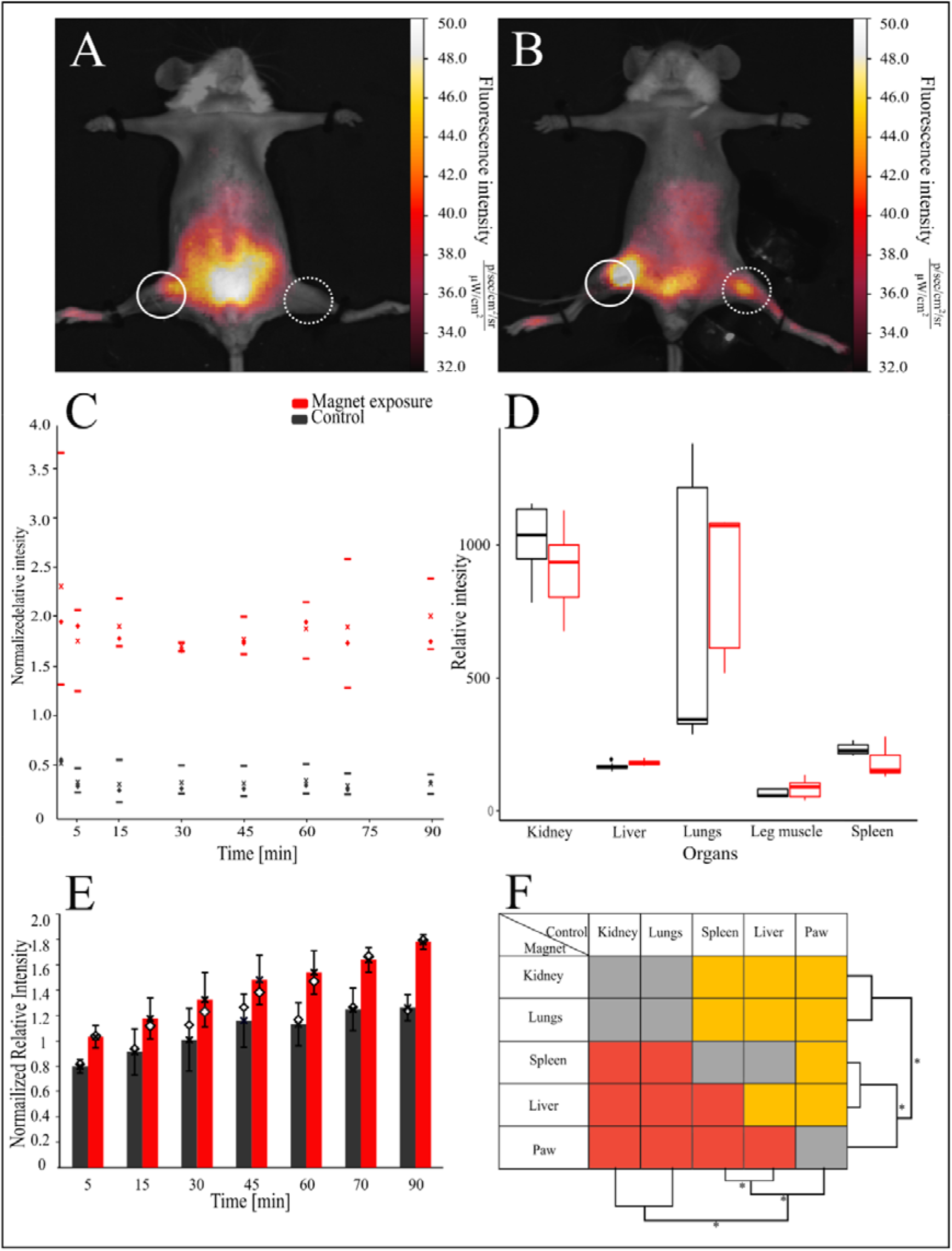
Distribution of the bacterial cells modified with magnetite nanoparticles and Cy-7 labeled PEI in the mice. The bacteria were detected in control mouse (A) and exposed under magnet mouse (B), respectively, dash circles indicate place of injection, solid circles – place of cell collecting. Relative fluorescence intensity of the signal in the paw under magnet exposure and normal distribution by blood flow was chosen to compare the accumulation of cells (C), where - Median, ⍰ - average, upper and lower borders are 25th to 75th percentile respectively. Normalized relative fluorescence was also assessed in the organs of animals (D). Introduction of bacteria led to increase of relative fluorescence intensities in mice bodies over time (E), where - Median, ⍰ - average and statistical significance of the distributed bacteria between organs (F), red – signal of the magnet group is lower than control, gray-not significant, orange – higher, respectively.

The fluorescent signal recorded from other parts of the body was increasing over time (Figure 2E). Throughout the whole time of the experiment we observed on average 1.3 times higher increase of the signal intensities from the body in the magnet group than in control (*P* < 0.05, see also **Supplementary Table 2**). Although the increment of signal in the group of animals exposed to the permanent magnet was observed, the signal intensity from the rest of the body was 1.8 times lower than the control at the end of the experiment (*P* < 0.001).

After animals were dissected we analyzed the fluorescent signals from the vital organs, which were 1.1 times significantly higher just in the liver within the magnet exposed group than in the non-exposed group of animals (*P* < 0.05), (Figure 2D). The results of paired statistical comparison between organs showed that the highest amount of the cells was accumulated in kidney and lungs (not significantly different, *P* > 0.05) with the following series of the reduction of the signal in liver>spleen>paw (*P* < 0.05 in inter pairs, Figure 2F).

## 3. Discussion

For the first time here we showed a proof of a new concept that a method of surface modification of bacterial cells by electrostatic deposition of PEs can increase the specificity of delivery of the EFBC to the site of an action by remote guiding more than 5 times (see Figure 2C). The specificity of our solution also exceeded more than 2 times over using microparticles of similar sizes, which could be attributed to a 4 times stronger permanent magnet used in the experiments.^[22]^ We expect that specificity of targeting could be improved even more since the deposited PEs have various amino or carboxyl groups that can enable adding specific receptors, proteins or antibodies that can target the tumor cells when bacteria are placed in the local environment of the cancers by the magnetic field.^[18,23,24]^ Since it was needed to gain a sufficient signal, it was injected a high dose of the EFBC (8 ⋅10^8^ CFU). As a result, there were many free circulating cells left, where we were able to capture only a small fraction of bacteria (up to 4.6%) in the magnet paw (see Figure 2 B), while the rest of the cells later passively accumulated in organs (see Figure 2 D). Therefore, if we want to increase the efficiency of targeting, the concentrations of cells should be lowered and optimized according to the power of the magnet source and vasculature capacity that would hold attached biomass. On a basis of safe concentrations known from literature that can be introduced to an organism, vasculature capacity and distribution of cells within the body determined as collected signals in our experiments, we can use at least 1000 times less bacteria to obtain similar results without extensive circulation of bacterial cells elsewhere in the body.^[2,25]^

Although in our experiments we did not use different modifications of the PE capsules, from the data obtained from the literature we can predict that the approach shown here can be further improved by using methods of modifications of the PE shell. One important condition using bacterial therapy is making bacteria stealth to the immune system. We can envision several mechanisms that would allow to gradually control the engagement of the immune system rather than obtaining a septic shock: (i) the PE capsules can hide bacterial membrane and toxic lipopolysaccharides (LPS) compounds from direct exposure, (ii) capsules can control the release of the cells as they create a strong barrier and (iii) specific coatings can increase stealth circulation. Accordingly, the cell wall of bacteria is composed of LPS, muramyl dipeptides and lipoteichoic acid, which are well known to be toxic components for the immune system.^[14]^ When the native bacteria are administered in high quantities, it is expected to have a strong inflammation with a high immune response followed by a septic shock. If we introduce native bacteria as high as 10^9^ CFU to BALB/c mice it should induce 50% lethality.^[25]^ However, although a high dose of bacteria (8 ⋅10^8^ CFU) was used, no lethal cases were observed during our experiments. We suppose that such a high viability can be a result of the presence of the PE shells, which could smoother the immune response of mice by shielding the LPS surface from direct contact with the immune cells. Specifically, the shielding was shown to be achieved for various inanimate PE capsules when they were introduced to an organism.^[26,27]^ Moreover, the potential systemic immune response could be prevented if we control the portion of the released EFBC that come in direct contact with the immune system. Based on our experiments performed *in vitro* (Figure 1E, 1F), it is shown that the bacteria stay alive being encapsulated. However, they need a longer time that is proportional to the capsule thickness to eventually escape. Therefore, introduction of bacteria with different thickness of the coating would enable gradual release of bacteria from shell and modulation of the intensiveness of the response of the immune system as only a particular portion of cells would release from the capsules at a certain time.

Eventually, the coated bacteria can be recognized by immunity via opsonization, which inevitably will engage activity of macrophages. Thus, to increase the stealth time of the capsules, specific coatings like polyethylene glycol (PEG) or polysaccharides could be used as they reduce hydrophobicity and surface charge, which are primal initiators of opsonization.^[28]^

Using the PE assembly can also help to overcome limitations of bacteria that lack targeting ability as it enables formation of the coatings responsive to remote guiding. In combination with controlled time of LPS introduction to the immune system and specific targeting, it can lead to a very high, but local immune response, which in return reduces the overall toxic effect. Besides, we expect that the cell growth will mostly occur in tumor hypoxic regions, since there are the most favorable growth conditions due to suppression of immunity.^[29,30]^ Bacteria can become even more potent by introduction of genetic elements for the expression of cytokines,^[29]^ cytotoxic agents ^[31]^ that additionally would contribute to fighting tumors or become more detectable by production of fluorescent dyes.^[32]^ However, we do not limit the application of this approach only to mitigate the cancer development as it allows us to use the coated bacteria as adjuvant systems of vaccines. Accordingly, if the parts of the bacterial cell envelopes can enhance the immune response,^[14,33]^ we can expect even higher elevation of the immunity if the whole bacterium is presented as a coated capsule but not exaggerating immune activity to engage the septic shock. In addition, the use of encapsulation can increase adherence of the probiotic bacteria ^[16]^ as well as their adjuvant activity.^[34,35]^

In our experiment we also analyzed the overall fluorescent intensities (Figure 2 E), which was unexpectedly gradually increasing during the duration of the experiment. This phenomenon we can speculate that occured at least due two reasons: (i) as a result of breaching of the shell integrity caused by either due to the bacterial cell division, or interaction with host organism, e.g. phagocytosis, eventually releasing labeled PEs or (ii) dissemination of bacterial cells through the body leading to saturation of the capillaries within the skin tissue that is covering the body surface resulting in a higher detectable signal. We observed a significant change of intensity after 15 min for the group of animals exposed to the magnet and 70 min for control (*P* < 0.01), which could not be attributed to the cell division as it would take at least 40 min under the best conditions to divide. Even, if the cells would be uptaken by the macrophages, killing of 40-70% of biomass (depends on type of cells and expression profile), it would take at least 2h with the following digestion and core, bacterial cell, decomposition for other 6 h.^[36]^ Also, we excluded the technical error, based on an improper set of the intensity scale, that might result in the signal increase. Hence, these discrepancies of different signal enhancement cannot be explained by shell degradation at least before the first division occurs. Here, we assume that the main reason could be related to a slower retention of saturation of the capillaries by the magnet compared to the control. From the literature it is known that the total surface area of capillaries (2000-27000 mm^2^) is much greater than the surface arterial vasculature (70 to 160 mm^2^) of mice.^[37]^ In contrast, blood volume in capillaries is much smaller in comparison to arteries and veins (5% against 15% and 64% respectively) as well as blood flow is not high.^[38]^ We suppose that in the long term, the number of freely circulating bacteria will be reduced by filling all capillaries and organs, depleting cells from blood of arteries and veins. The cells will increase the detectable surface signal being stuck near the skin surface region. Since the capillaries are highly branched, small differences in the concentration of the cells retained by the magnet, eventually might cause slower saturation of capillaries, till the magnet is removed and held leftovers of bacteria further fill the rest of capillaries. As we did not prolong the experiment, some amount of the cells could be still held in place of the magnet, not reaching control values and as a result the overall intensity was much lower. In addition, retention of the cells by the magnet over time might cause formation of the soft aggregates, which will not be able to fill capillaries due to an excessive size. Eventually, at the end of the experiment, the cells were mainly distributed in kidney and lungs, similar as reported for uncoated bacteria^[30]^ or silica particles of micron size.^[39]^ From literature it is also known that nanosized particles tend to be accumulated in spleen, lungs, liver and kidney.^[40–42]^ Since the vascular volumes of the organs as well as blood flow have the highest values in lungs and kidney,^[43]^ it could explain the fact that bacteria were mostly accumulated in these organs with the following reduction to liver, spleen and paw. Therefore, for successful delivery, it has to be taken into account that difference in dissemination from this perspective can be considered as vasculature, blood flow as well as size dependent.

## 4. Conclusions

Summing up, we designed a solution for improving bacterial therapy by using the method of electrostatic deposition. We prepared the magnetically responsive bacterial cells, which were detectable in the far red spectrum. Using the permanent magnet allowed us to increase the specificity of targeting more than 5 times compared to the control group. It was found that introduction of a high dose of the coated cells did not lead to animal death throughout the whole experiment. As the polyelectrolyte layers were deposited on the cells, it resulted in the controlled time of the cell division, which might be used for gradual introduction of the coated bacteria to the immunity. Such an approach of encapsulation opens new possibilities for using and controlling various microorganisms and makes new concepts in drug delivery as well as in preparation of adjuvants in vaccines.

## 5. Materials and methods

### 5.1 Bacteria strains and growth conditions

In our experimental simulation of targeted bacterial therapy we used cells of non motile *Escherichia coli* TOP 10 strain (F– mcrA Δ(mrr-hsdRMS-mcrBC)Φ80lacZΔM15 ΔlacX74 recA1 araD139 Δ(ara leu) 7697 galU galK rpsL (StrR) endA1 nupG), containing pRSET-emGFP plasmid. The plasmid contains the T7 promoter region upstream of the emGFP reporter gene and ApR cassette, which enabled us to observe cells by confocal microscope and to control bacterial contaminants by amending ampicillin in media, respectively. The transformants were cultivated at 37 □ on nutrient agar plates (Sigma-Aldrich) supplemented with ampicillin (100µg mL^-1^, Sigma-Aldrich) (NAamp). From the overnight culture, we transferred 1 mL into the 100 mL fresh medium and incubated until obtaining the optical densities appropriate for conducting the particular experiments. All liquid cultures were incubated by shaking on a rotary shaker at 37 □ and 150 rpm.

### 5.2 Preparation of PEs for layer by layer encapsulation

We used the negatively charged sodium poly(styrene sulfonate) (PSS, MW=70 kDa) and the positively charged poly(ethyleneimine) PEs (PEI, MW=600 kDa), both purchased from Sigma-Aldrich, for cell encapsulation based on the electrostatic principle.^[44]^ The solutions of PEs in MQ water (2.5mg mL^-1^, pH 7 adjusted by NaOH or HCl) were prepared by solubilizing PEs firstly by stirring and then by the sonication (35kHz, 100W, 15min). PEI was stained with Cyanine7 (Cy7, Lumiprobe, USA) according to protocol of NHS ester labeling of amino biomolecules.^[45]^ Briefly, the Cy7 (9 mg in 9 mL of DMSO) solution was added to PEI solution (2.5 mg mL^-1^ in 40 mL of water with pH 8.4) in 50 mL tube and incubated for 4 h at room temperature under constant stirring conditions. To remove the residual dye after labeling the solution was dialyzed for 3 days using a dialysis tube (Orange Scientific) with nominal molecular weight limits between 12 and 14 kDa and titrated up to pH 7. All the PEs were sterilized by filtration using 0.2 µm sterile filters.

### 5.3 Encapsulation procedure

Prior to the procedure of encapsulation of cells in PE layers, bacterial cells were grown at 37 □ by shaking at 150rpm until reaching OD_600_ = 0.2 measured in a 200 µl volume on a microtiter plate. The obtained culture of such a densities of bacterial cells were further concentrated by centrifugation of the 50 mL of bacterial culture at 5000 g for 6 min and then washed three times by the centrifugation (5000g for 5 min) and finally resuspended in 30 mL of 0.9 % NaCl. The cells were coated in layers of PEI/PSS/PEI/magnetite-NP/(PEI/PSS)_2_ by the following procedure. Firstly, positively charged PEI layer was deposited by adding 0.25% solution of PEI in MQ water (pH=7, adjusted with HCl) to the washed and concentrated cells of final OD_600_=1.2 in 1:1 v/v ratio with PEI and incubated at room temperature for 5 minutes. Unattached PEI was washed out from the suspension by centrifugation at 900 g for 2 min. The obtained pellet was 2 times washed by the gentle pouring of 1mL of 0.9% NaCl not resuspending the pellet. The PEI coated cells were washed in 0.9% NaCl solution and then the second negatively charged PSS layer was deposited (pH 7 adjusted with NaOH) using the same procedure as for PEI except the cells were centrifuged at 1500 g for 3 min. After addition of another PEI layer the layer of magnetite NPs was added. The magnetite NPs (12 nm average size), were synthesized according to German et al. (2013).^[46]^ The NPs were incubated with the cells for 5 minutes (1 mg mL^-1^ particles, 1:1 v/v particles to cells). The excess of magnetite particles was washed out by centrifugation at 1500 g for 3 min. The coated cells were collected by the magnet, additionally washed 2 times by pipetting and then resuspended in 0.9% NaCl solution. After magnetite, the layers of PEI-Cy 7/PSS were deposited in the same way as there were deposited in the first layers of PE on the surface of the bacteria. Finally, we obtained 8 layers of the PEs on the bacterial cells.

### 5.4 SEM and EDS analysis

For obtaining the best possible images using SEM visualization, samples were prepared using different protocols: (i) air drying at room temperature without any fixatives, (ii) the freeze drying procedure and (iii) chemical fixation with drying at 40 □. For normal drying the samples of the coated and control cells (intact, free of magnetic NPs) were suspended in MQ water and placed on silicon wafers to be dried at room temperature for 72 hours. For the freeze drying, the cells were resuspended in MQ and pre-frozen in liquid nitrogen. Further, the samples were put in the freeze dryer (Christ Gamma 1-16 LSC Plus) and left for 72 h at -50 □. For chemical fixation the cells were prepared by the following procedure.^[47]^ Briefly, the cells were fixed with 0.5% glutaraldehyde and 1% formaldehyde in a 0.1 M cocodylate buffer 0.1 M and transferred on a clean glass. After washing with cocodylate, 1% osmium tetroxide was added and left for 1 h. After all, samples were rinsed with water and incubated for 5 min in gradient dilutions of ethanol (30%, 50%, 70%, 96%), replaced with hexamethyldisilazane and left for drying overnight at 40 □. All the samples from different protocols were mounted on SEM stubs, platinum-sputtered and observed in high vacuum SEM (Jeol JSM-7600F) with a field emission gun, at low voltages, to observe the structure of the cell. For EDS silicon drift detector (SDD) from Oxford (S Max, 20mm^2^) was used. To excite the Kα a higher excitation voltage was applied.

### 5.5 Time-lapse confocal microscopy (TLCM)

To enable the precise measurement using the TLCM approach we needed to distribute cells on a thin planar surface and provide them with appropriate growth conditions for a few hours of observations. Accordingly, we prepared a chamber for the cell incubation as described previously.^[21,48]^ Briefly, the nutrient agar was poured inside a cut of 1cm x 3 cm of a double-sided tape attached to a clean glass. Before solidifying the cover slip glass was carefully deposited to make a smooth NA layer. The chamber was left for 10 min 4 □ to form a solid NA matrix. The non attached slide glass was then removed, the protective layer of the double sided tape was pilled off and then 4 µl of culture (final OD_600_ approx. 0.4) was evenly distributed over the exposed NA surface. At the end we sealed the chamber with the cover slip by attaching it on the sticky surface of the frame enclosing the NA.

The chamber with the cells was equilibrated at 37 □ for 15 min and transferred to the confocal microscope (Leica TCS SP8X confocal laser scanning microscope equipped with temperature control system cube and box) that was thermostated at 37 □. The TLCM was performed at 1000x magnification using an objective lens (HCX PL APO 100x/1.44 OIL) immersed in immersion oil. We observed cell growth and division using the excitation at 489, emissions 525/25 and 750 842/33 with 850-900V and 750-800V gains for emGFP and Cy7 fluorescence, respectively. The morphology of cells and amount of non-fluorescent ones was observed using white light and condenser as the objective lens.

### 5.6 Analysis of the TLCM micrographs

To determine the growth properties of the entrapped cells in PE layers we analyzed images using Fiji software.^[49]^ In prior to the experiment we firstly selected fields under the microscope occupied by similar amounts of cells. From the obtained fluorescent images forming hyperstack we deleted those that were of low contrast or out of the focus. The prepared hyperstacks of the images were then converted to the stack of binary images using the “make binary” plugin. On such prepared images we measured the surface area of cells at the consecutive time points by using the “analyze particles” plugin to determine temporal changes of the cell biomass. The experiments were performed in triplicates and the t-test was used for statistical comparison of the amount of cell biomass per time per group of cells with different numbers of PE layers.

### 5.7 In vivo biodistribution

All experiments were performed according to the relevant institutional (National Research Ogarev Mordovia State University, Russia) and international regulations of the Geneva Convention of 1985 (International Guiding Principles for Biomedical Research Involving Animals). Animal ethics clearance was approved by the ethical committee (protocol № 50 from 29.05.2017). The biodistribution of the encapsulated bacterial cells was investigated in 2 groups of 3 BALB/c female mice 6-8 weeks old with weight distribution 20–25 g. The intra-arterial injections of suspension of bacteria were performed on immobilized anesthetized mice. For general anesthesia Zoletil (60 mg kg^-1^, Virbac SA, Carros, France) and Rometar (and 10 mg kg^-1^, Spofa, Czech Republic) mixture was used via intraperitoneal injection. The intra-arterial injection was performed through the polyethylene catheter (PE-10 tip, Scientific Commodities INC., Lake Havasu City, Arizona), which was introduced into the right femoral artery. During the implantation the catheter was filled with isotonic sodium chloride solution. The bacterial cells were resuspended in sodium chloride solution of approximate OD_600_ = 1 (measured in a 200 µl of microtiter plate). The volume of bacterial suspension administered in the mouse was 200µl.

For all animals, the bacterial cells were injected in the right femoral artery and collected in the left paw by placing the permanent magnet NdFeB (1400 mT remnant magnetization, diameter -50 mm, height -20 mm) for 60 min, which was then removed. The control group of animals was treated in the same way except permanent magnet was not used.

### 5.8 In Vivo fluorescent imaging

We measured distribution of Cy7 labeled bacteria using IVIS® Lumina imaging system (Xenogen Corp.) with a filter set (excitation, 710-760 nm; emission, 810-875 nm). All the fluorescence images were acquired with 0.2 s exposure. The intensities of fluorescent signals on the obtained pictures were also analyzed in Fiji software.^[49]^ All images were converted to the gray-scale signal using the “RGB to luminance” plugin. Prior to analysis, all the signals were normalized according to the initial base level. The gray-scaled signals from the paw were normalized to the signals from the abdominal area and intensities were measured over the area with the resolution of 400 pxl. Intensities of organs were measured for the whole area of a particular one and normalized by the weight of a single organ. Absolute intensity of every mouse was measured according to the same procedure, except the rectangular frame was used to calculate change in fluorescence. All experiments were performed using 3 animals per tested group. Nonparametric Mann Whitney two-tailed and Kruskal-Wallis tests were used for statistical comparison of signals obtained from different body parts as well as from different organs when animals were sacrificed and dissected.

## Acknowledgements

Work was supported by Slovenian-Russian bilateral project (BI-RU/16-18-039), Slovenian national projects (J4-7640, J1-6746, J3-1762, J1-9194, J7-9400, and P1-0143), Flemish-Slovenian research project: Bioavailable mercury methylation in the Adriatic sea (BE MERMAiD, grant agreement N1-0100), Helmholtz Association (grant PIE-0007 CROSSING), European Urban Initiative Actions founded project Applause (Grant agreement UIA02-228), 2019 -2023 (EU -Horizon 2020): InteGRated systems for Effective ENvironmEntal Remediation (GREENER, grant agreement 826312) and European Commission (SurfBio project, grant no.: 952379).

