## Supplemental_tables_1_2 for "*In vivo* capture of bacterial cells by remote guiding"

By *in vivo* imaging of the electrostatically fictionalized bacterial cells we revealed: (i) concentration of the cells can be locally increased more 5 times by using the magnet (ii) the exposed place has higher intensity values even when the magnet removed (iii) different exposure type affects the cell distribution and (iv) polyelectrolytes can regulate growth rate of the bacteria.

Iaroslav Rybkin^1,2,3,4^, Sergey Pinyaev^5^, Olga Sindeeva^3,6^, Sergey German^3,6^, Maja Koblar^2,4^, Nikolay Pyataev^5^, Miran Čeh^2^, Dmitry Gorin^3,6^, Gleb Sukhorukov^3,7^, Aleš Lapanje^2^*

Title *In vivo* capture of bacterial cells by remote guiding

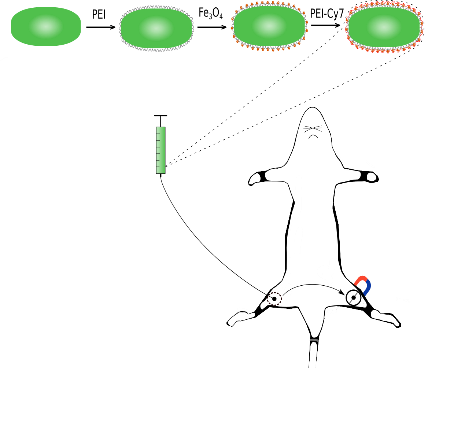

Supporting Information

*In vivo* capture of bacterial cells by remote guiding

Iaroslav Rybkin^1,2,3,4^, Sergey Pinyaev^5^, Olga Sindeeva^3,6^, Sergey German^3,6^, Maja Koblar^2,4^, Nikolay Pyataev^5^, Miran Čeh^2^, Dmitry Gorin^3,6^, Gleb Sukhorukov^3,7^, Aleš Lapanje^2^*

Graphical abstract

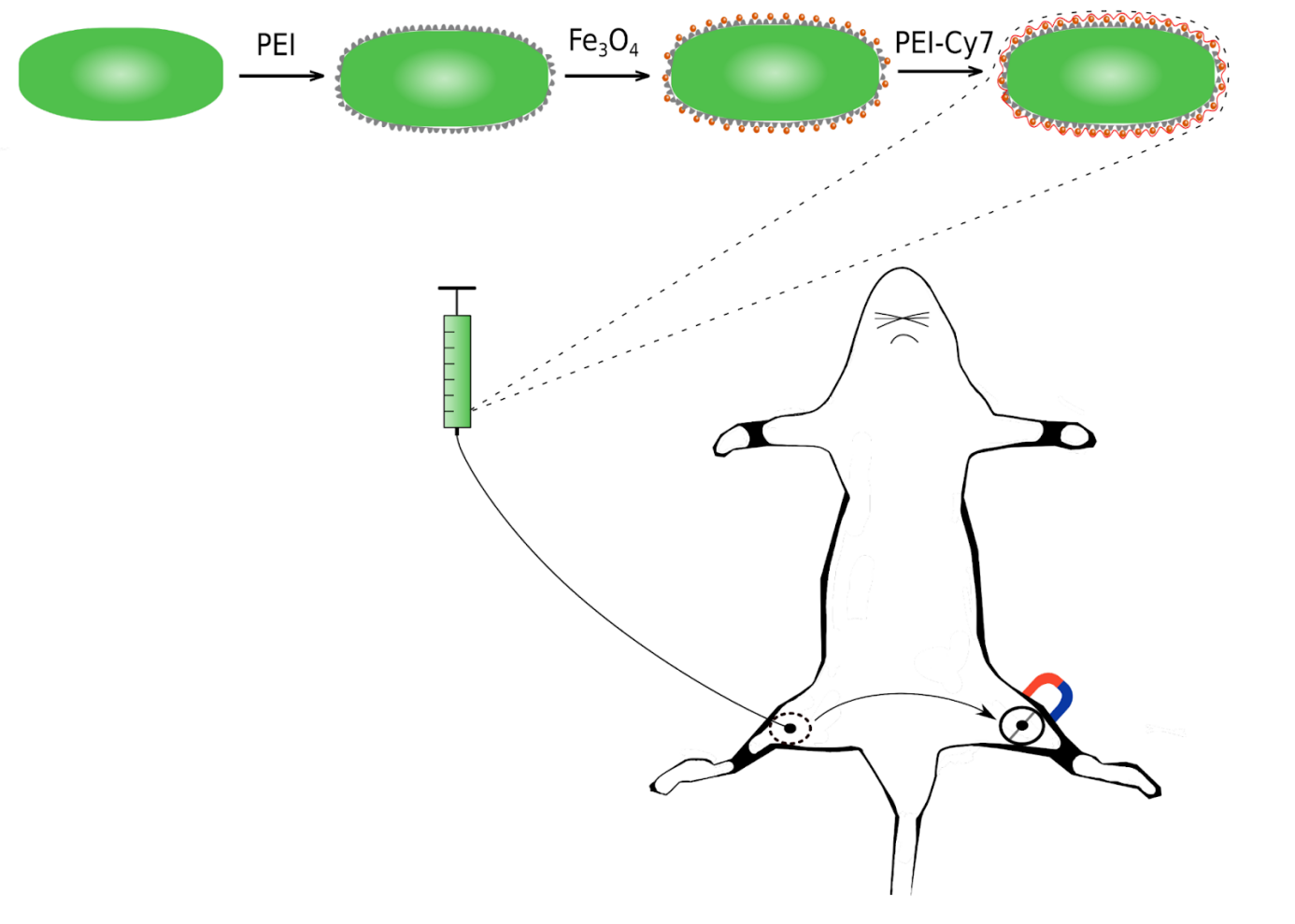

**Supplementary Table 1.**  EDS spectrum of the coated and non-coated by magnetite bacterial cells

| Spectrum | C % | O % | Na % | Si % | Fe % | Cl % | Pt % |
| --- | --- | --- | --- | --- | --- | --- | --- |
| Spectrum 1 control | 36 | 2.9 | N/A | 60.8 | N/A | 0.1 | 0.2 |
| Spectrum 2 control | 51 | 4.7 | 0.1 | 44.1 | N/A | N/A | 0.2 |
| Spectrum 3 control | 45.1 | 4.3 | 0.1 | 50.3 | N/A | 0.1 | N/A |
| Spectrum 4 control | 35.6 | 2.8 | N/A | 61.4 | N/A | N/A | 0.2 |
| Spectrum 5 control | 37.3 | 2.7 | 0.1 | 59.6 | N/A | N/A | 0.2 |
| Spectrum 1 coated cells | 41.3 | 9 | 1 | 47.6 | 0.6 | 0.3 | 0.2 |
| Spectrum 2 coated cells | 44.6 | 10.3 | 0.6 | 43.9 | 0.3 | 0.3 | 0.1 |
| Spectrum 3 coated cells | 41.3 | 7.7 | 0.4 | 50.1 | 0.1 | 0.3 | 0.2 |
| Spectrum 4 coated cells | 40.6 | 6 | 0.3 | 52.6 | 0.1 | 0.2 | 0.2 |
| Spectrum 5 coated cells | 54.3 | 12.9 | 1 | 31.2 | 0.2 | 0.3 | 0.2 |

**Supplementary Table 2.**  Modes of relative increased intensities when fluorescently labeled bacteria are introduced in the body of mice

| Samples | Time [min] | Mode 1 | Mode 2 | Mode 3 |
| --- | --- | --- | --- | --- |
| Control mice | 5 | 0.8 | NA | NA |
|  | 15 | 0.7 | 1.1 | 0.9 |
|  | 30 | 0.7 | 1.2 | 1.1 |
|  | 45 | 0.9 | 1.3 | NA |
|  | 60 | 0.9 | NA | NA |
|  | 70 | 1.3 | NA | NA |
|  | 90 | 1.2 | NA | NA |
| Magnet exposed mice | 5 | 1 | NA | NA |
|  | 15 | 1.1 | 1.4 | NA |
|  | 30 | 1.2 | NA | NA |
|  | 45 | 1.4 | NA | NA |
|  | 60 | 1.4 | NA | NA |
|  | 70 | 1.7 | 1.6 | NA |
|  | 90 | 1.8 | NA | NA |
